# Positive selection on mitochondria may eliminate heritable microbes from arthropod populations

**DOI:** 10.1101/2021.03.26.437186

**Authors:** Andy Fenton, M Florencia Camus, Gregory D D Hurst

## Abstract

The majority of arthropod species carry facultative heritable microbes, bacteria that are passed from mother to offspring, and which may contribute to host function. These symbionts are coinherited down the maternal line with mitochondria, and selection favouring either new symbionts, or new symbiont variants, is known to drive loss of mitochondrial diversity as a correlated response. More recently, evidence has accumulated of episodic directional selection on mitochondria. We therefore examined the reciprocal interaction and model the impact of selection on mitochondrial DNA (mtDNA) on symbiont frequency. We performed this for three generic scenarios: a fixed benefit to the host carrying the symbiont, a benefit that decreased with symbiont frequency, and a benefit that increased with symbiont frequency. We find that direct selection on mtDNA can drive symbionts out of the population under some circumstances. Symbiont extinction occurs where the positively selected mtDNA mutation occurs initially in an individual that is uninfected with the symbiont, and the symbiont is initially at low frequency. When, in contrast, the positively selected mtDNA mutation occurs in a symbiont infected individual, the mutation becomes fixed and in doing so removes symbiont variation from the population. Given low frequency symbiont infections are common in natural populations, and selection on mtDNA is also considered to occur frequently, we conclude that mtDNA driven loss of symbionts represents a novel mechanism driving loss of facultative heritable microbes. We conclude further that the molecular evolution of symbionts and mitochondria, which has previously been viewed from a perspective of selection on symbionts driving the evolution of a neutral mtDNA marker, should be reappraised in the light of positive selection on mtDNA. Where low mtDNA and symbiont genetic diversity are observed, it should not be assumed to be a consequences of selection acting on the symbiont.

## Introduction

The majority of arthropod species carry heritable microbes – bacteria, viruses and fungi that pass from a female to her progeny (1). These microbes modify the biology of their host individual, acting both as beneficial symbionts that contribute to organismal function, and as reproductive parasites. Contributions to function commonly include synthesis of amino acids and essential cofactors and protection of the host against attack by natural enemies (virus, parasitic wasps, nematode worms, fungi, predators) (2). Reproductive parasitism is a consequence of the exclusive maternal inheritance of the microbe, which selects for strains that bias investment towards the production and survival of female offspring over male, and for incompatibility between infected males and uninfected females (3). These individual impacts affect population and community dynamics and are additionally important drivers of host evolution.

Heritable microbes are maternally inherited alongside mitochondria, and it has long been recognised that the spread of heritable microbes leads to the spread of the mitotype originally associated with the symbiont (4). Thus, mitochondrial evolution and diversity is a product in part of the recency with which either a new symbiont, or a mutation of an existing symbiont strain, has spread into the population (5). This process is sufficiently strong that symbionts may also drive mitochondrial introgression across species boundaries following rare hybridization events (6), and as such represent potential disruptors of the ‘barcoding gap’ (7).

More recently, it has become clear that mitochondria are not the neutral marker they have been historically considered but are in fact themselves subject to direct selection (7–9). Direct selection may be associated with their vital role in energy and heat production, as well as dietary and thermal adaptation (10,11). Mitonuclear coadaptation is also recognised, in which selection acts jointly on mitochondrial and nuclear variants to maintain a functional integrated mito-nuclear system (12, 13). It is thus clear that the current view – that mitochondrial diversity is shaped by symbionts – is likely to represent one component of a reciprocal interaction. We would additionally expect selection on mitochondria to impact on the dynamics and diversity of heritable microbes. If this is true, the outcomes of mitochondrial selection may vary depending on the phenotype of, and direct selection pressures experienced by, the symbiont. However, these possibilities have yet to be formally explored.

In this paper, we use mathematical models to explore the impact of positive selection on mitochondria upon the dynamics of facultative heritable symbionts. We perform this analysis for three dynamical categories of heritable microbe. First, we consider symbionts maintained by a balance of a direct drive phenotype that is fixed in magnitude, alongside segregational loss. This model class approximates the dynamics of both reproductive parasitic phenotypes like male-killing (14), and nutritional benefits. We then consider symbionts maintained by a benefit that declines as they become more common (negative frequency dependence). This class of models is applicable to defensive symbionts where symbiont spread is driven by imparting resistance to natural enemies, but as the symbiont becomes more frequent, so natural enemy attack rates (and the benefits of symbiont carriage) may decline (15). Finally, we explore symbionts where the symbiont drive is positively frequency dependent, that is to say the advantage of having the symbiont becomes stronger when the symbiont is more common. For this class, we model the particular case of cytoplasmic incompatibility, where the uninfected zygotes die if the fertilizing sperm comes from an infected male (16).

## Modelling framework

To understand the coevolutionary dynamics of a potential new mitotype and a symbiont, we initially assume the symbiont is at equilibrium with the resident mitotype and seek the criteria which leads to invasion (and possible replacement of the resident mitotype) by a new mitotype, alongside the consequences for the persistence and equilibrium prevalence of the symbiont. In each case, we analyse the dynamics of a novel beneficial mitotype introduced into symbiont infected, or symbiont uninfected, individuals separately. All mathematical details are described in the Appendix. The three frameworks for symbiont dynamics are modelled generically as either a fixed benefit, a negative frequency dependent benefit, or CI with a positive frequency benefit. For ease, these formulations are simplified from real world dynamics (e.g. the fixed benefit to male-killing would be diluted by cannibalism of dead males by uninfected sibling females generated through segregation; negative frequency dependent selection approximation does not include eco-evolutionary dynamics: (15)); thus they approximate real world scenarios without precisely mirroring them.

For all scenarios, we model the change in frequencies of individuals that carry the resident mitotype and the symbiont (*p*), those that carry the resident mitotype but not the symbiont (*q*), those that carry the new mitotype and the symbiont (*r*) and those carrying the new mitotype but not the symbiont (*s*). We initially assume the symbiont is at equilibrium with the resident mitotype (i.e., the system is at 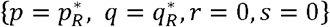, where 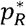 and 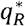 are the relevant equilibrium frequencies in the presence of the resident mitotype alone). A new mitotype then emerges that confers a fitness benefit of *t* to all individuals that carry it, exploring values up to *t* =0.02, reflecting strong selection on the mtDNA. This novel mitotype may emerge in either a symbiont-infected individual or one without the symbiont, and these initial conditions are examined separately. We then explore the stability criteria to establish the criteria for this invading mitotype to establish, and the consequences for persistence and prevalence of the symbiont. All simulations were carried out using Mathematica v12.1 (code available at https://doi.org/10.6084/m9.figshare.c.5354999.v1).

## Results

### i) Facultative heritable symbionts with a fixed benefit

In our mean field model, the symbiont is present in the population if the rate of symbiont’s segregational loss, *μ*, is less than *B*/(1 + *B*), where B is the net benefit of symbiont infection (see Appendix i). Positively selected mitochondria (*t*>0) that arise in a symbiont-infected individual spread to fixation. Symbiont prevalence increases during spread, as the advantageous mitotype is initially more common within symbiont infected individuals than uninfected ones. However, the prevalence of the symbiont at equilibrium is not altered, as the beneficial mitotype later establishes into the uninfected population following symbiont segregational loss (Figure 1). This process is easily visualised as generating a novel cytotype (symbiont + advantageous mitotype) that has higher benefit than all others. Its spread displaces other symbiont strains from the population (no additional mtDNA benefit), and segregational loss of the symbiont results in the mtDNA also invading through the uninfected portion of the population.

**Figure 1:**
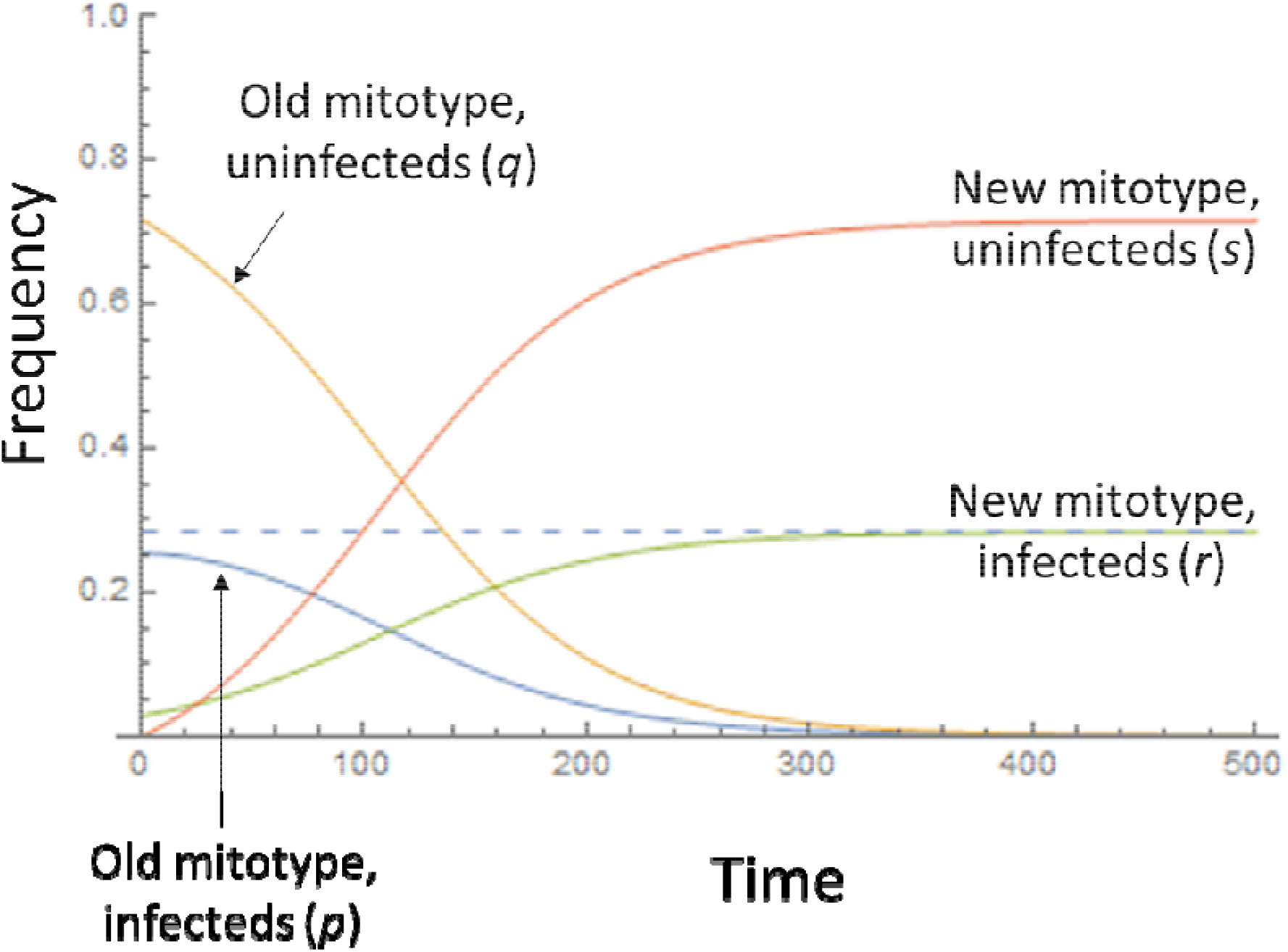
Dynamics of a novel advantageous mitochondria (*t* = 0.02) arising in a symbiont-infected individual in a population initially at equilibrium with that symbiont. Blue line – frequency of symbiont with ancestral mitotype; orange line – frequency of uninfected females with ancestral mitotype; green line–frequency of symbiont-infected individual with novel beneficial mitotype; red line – frequency of uninfected females with novel beneficial mitotype. Other parameters: *B* = 0.075; μ = 0.05.

A different pattern is observed when the novel beneficial mitotype mutation arises originally in an uninfected individual (Figure 2; see Appendix i for mathematical details). When a threshold benefit to the novel mitotype of *t* is reached (above the diagonal line given by *t* = *B*(1 – *μ*) – *μ*), the novel mitotype-infected females leave a greater number of progeny than the symbiont-infected females leave symbiont-infected progeny, and this mitotype invades and displaces the heritable microbe from the population. If this threshold is not reached (*t* < *B*(1 – *μ*) – *μ*; below the diagonal line in Fig 2), then the novel mitotype does not invade if it arises in an uninfected individual, despite being advantageous compared to the ancestor. For realistic values of mitotype advantage (*t* = 0.02, or a 2% benefit to the novel mitotype), exclusion is confined to symbionts that exist at low prevalence (these either have low benefit, or a higher benefit with high rates of segregational loss).

**Figure 2:**
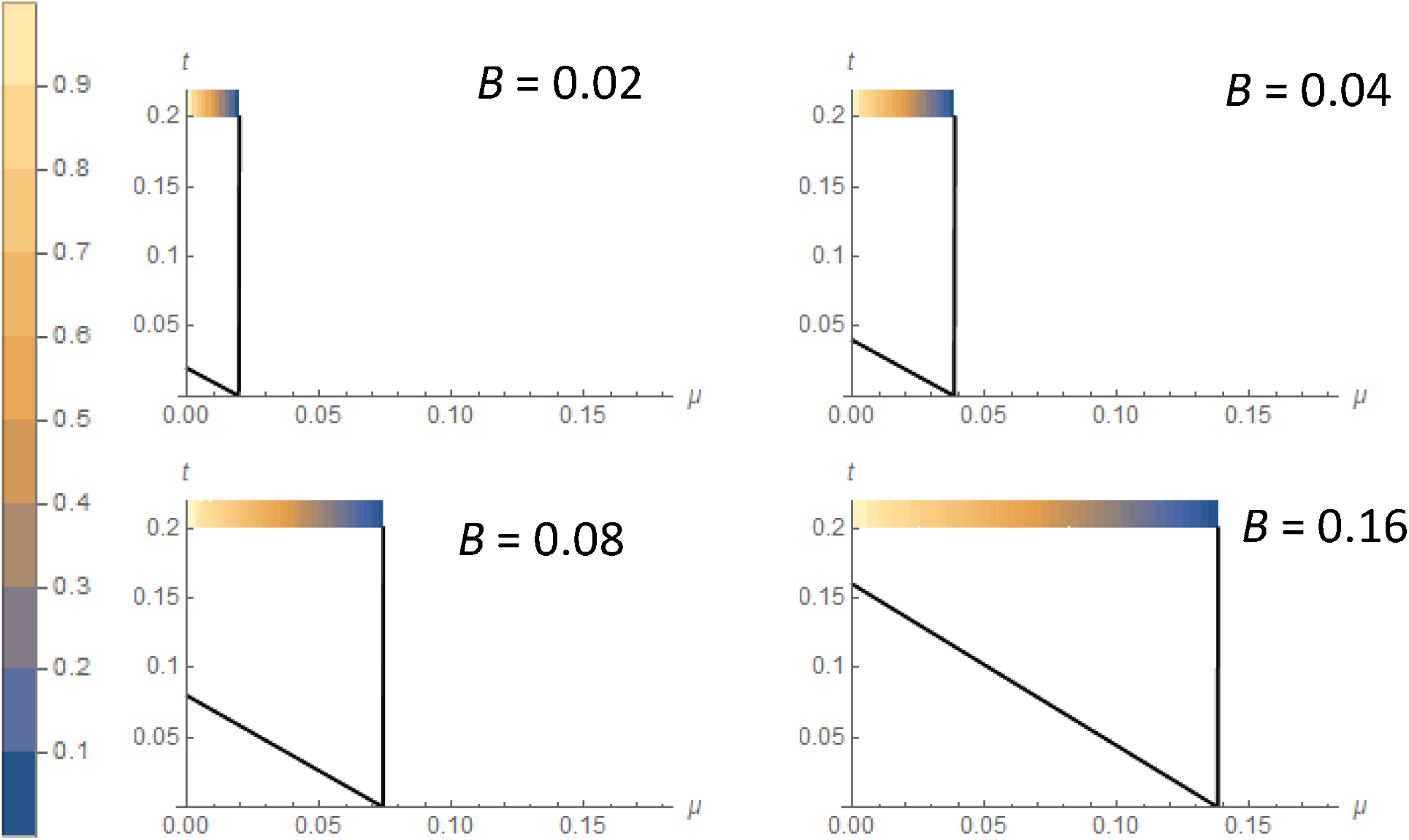
Conditions for invasion of a novel beneficial mitotype arising in a symbiont-uninfected individual where the symbiont has direct benefit to its host *b*. X axis represent rate of segregational loss (μ), and the heatmap the frequency of the symbiont at equilibrium, before mitotype invasion. Y axis represents the benefit of the novel mitotype (*t*). Spread and fixation of the novel mitotype occurs above the diagonal line. The symbiont cannot persist to the right of the vertical line given by _______.

### ii) Facultative heritable symbionts whose benefit declines as they become more common

We modelled the dynamics of mtDNA for a symbiont whose benefit declined with its overall frequency in the population, for example to mimic a defensive symbiont. For these symbionts, the frequency dependent benefit exists alongside a fixed cost of carrying the symbiont *c*, such that there is a polymorphic equilibrium for symbiont infection, at a frequency where the benefit of the phenotype is counterbalanced by the cost of infection and any segregational loss 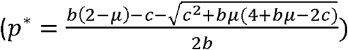. Part of our analysis focused in detail on the special case where a symbiont infected/uninfected polymorphism is retained without segregation (*μ* = 0) (a unique feature of negative frequency dependent advantage models for mean field formulations; for all other classes of mean field model, net beneficial symbionts fix if *μ* = 0).

A novel advantageous mitotype arising in an infected individual will spread through to fixation and, if there is any level of symbiont segregational loss, symbiont equilibrium prevalence will be unaltered as segregational loss results in the beneficial mitotype flowing into the uninfected fraction of the population (see Appendix ii). This process will also lead to the loss of symbiont diversity.

For the special case of no segregational loss of the symbiont (μ = 0), a novel advantageous mitotype in the infected portion of the population invades that section of the population but does not pass across to the uninfected portion by segregation. This portion thus retains the ancestral mitotype. If the mitotype is sufficiently advantageous to overcome fixed costs of symbiont infection (*t* > (*b* – *c*)), the symbiont becomes beneficial at all frequencies, at which point the symbiont will fix. Otherwise, the symbiont-infected portion increases in frequency associated with the additional benefit of carrying the new mitotype, and the ancestral mitotype persists in the symbiont uninfected fraction of the population, which declines in frequency. Mitotype variation becomes partitioned between symbiont-bearing and symbiont uninfected pools.

We then examined the dynamics of a novel advantageous mitotype arising in the symbiont-uninfected portion of the population (Figure 3). The conditions for the invasion of this mitotype are less restrictive than for a fixed benefit symbiont, as the negative frequency dependence of symbiont impacts makes the symbiont-infected type only weakly beneficial at the point of invasion of the novel mitotype. At the point of invasion, the symbiont is maintained polymorphic by a benefit – segregational loss balance, and mitotype invasion thus occurs when the benefit of the novel mitotype overcomes the rate of segregational loss from the symbiont infected section at equilibrium (above the lower curve in Figure 3). The invasion of this mitotype lowers symbiont prevalence. The ultimate fate of the mitotype then depends on its level of benefit. Above the diagonal line in Figure 3 (i.e., *t* > (1 + *b* – *c*)(1 – *μ*) − 1; see Appendix ii), the benefit of the novel mitotype exceeds that of the symbiont at low symbiont frequency, and the symbiont is displaced. In the area between the two lines, the novel mitotype invades the population but does not displace the symbiont. Here, females carrying the novel mitotype leave more progeny than symbiont-infected females when the symbiont is initially at equilibrium, allowing it to invade, but this fitness gap narrows as the symbiont declines in frequency and its benefit increases. These conditions create an equilibrium at which the symbiont-infected type has the ancestral mitotype, and the symbiont uninfected portion carries the novel mitotype (though not at fixation in this class, due to segregational loss of the symbiont dripping the mitotype from the infected compartment across to the uninfected compartment) (Figure 4).

**Figure 3:**
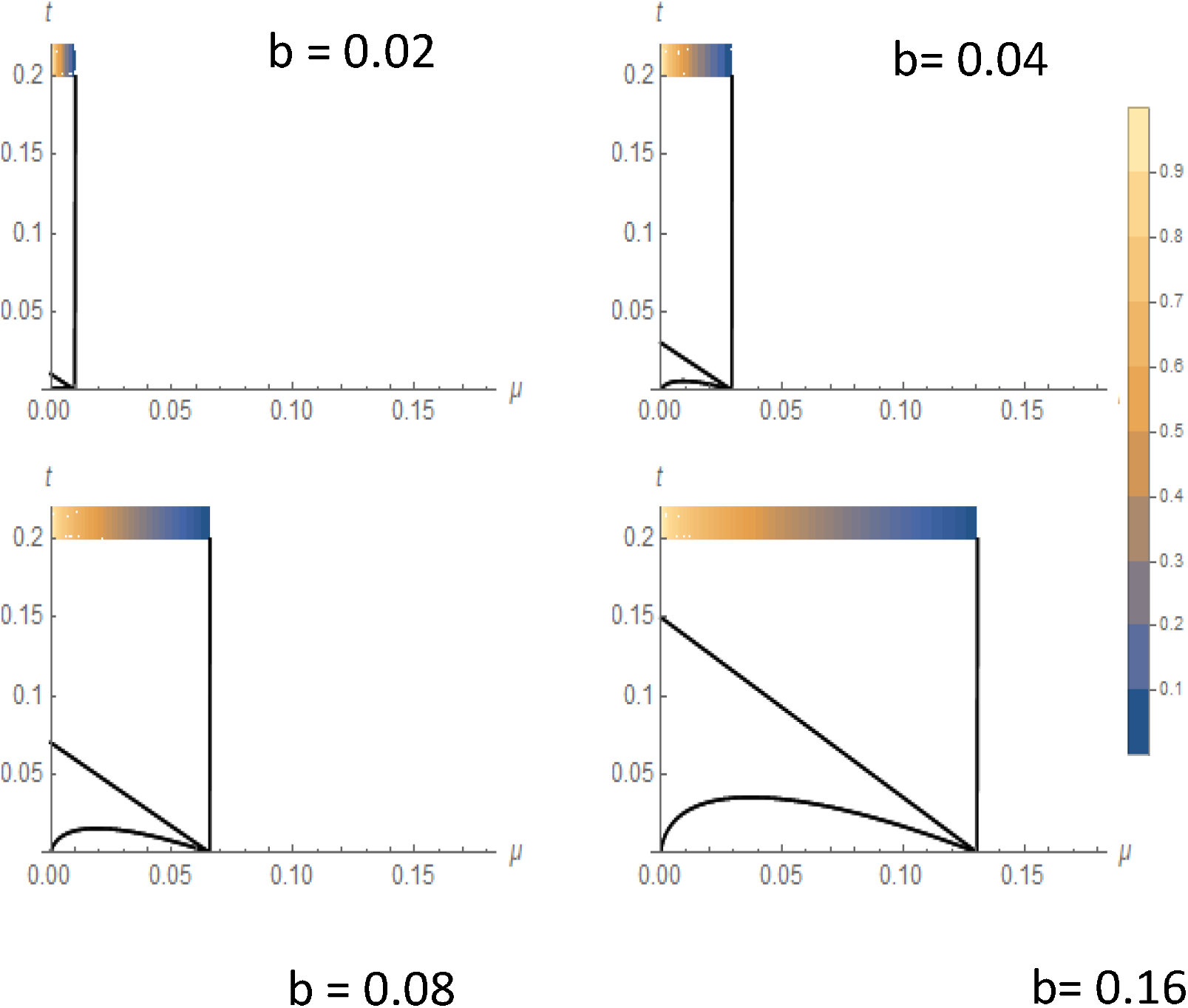
Conditions for invasion of a novel beneficial mitotype arising in a symbiont uninfected individual where the symbiont has direct benefit to its host b that declines with frequency of the symbiont. X axis is the rate of segregational loss (μ), and the heatmap the frequency of the symbiont at equilibrium, before mitotype invasion. Y axis represents the benefit of the novel mitotype (*t*). Spread and fixation of the novel mitotype occurs above the diagonal line; beneath this line but above the curve is an area where the mitotype invades but does not fix, remaining polymorphic within uninfected individuals. The symbiont cannot persist to the right of the vertical line given by ________. The cost of symbiont carriage *c* = 0.05.

**Figure 4:**
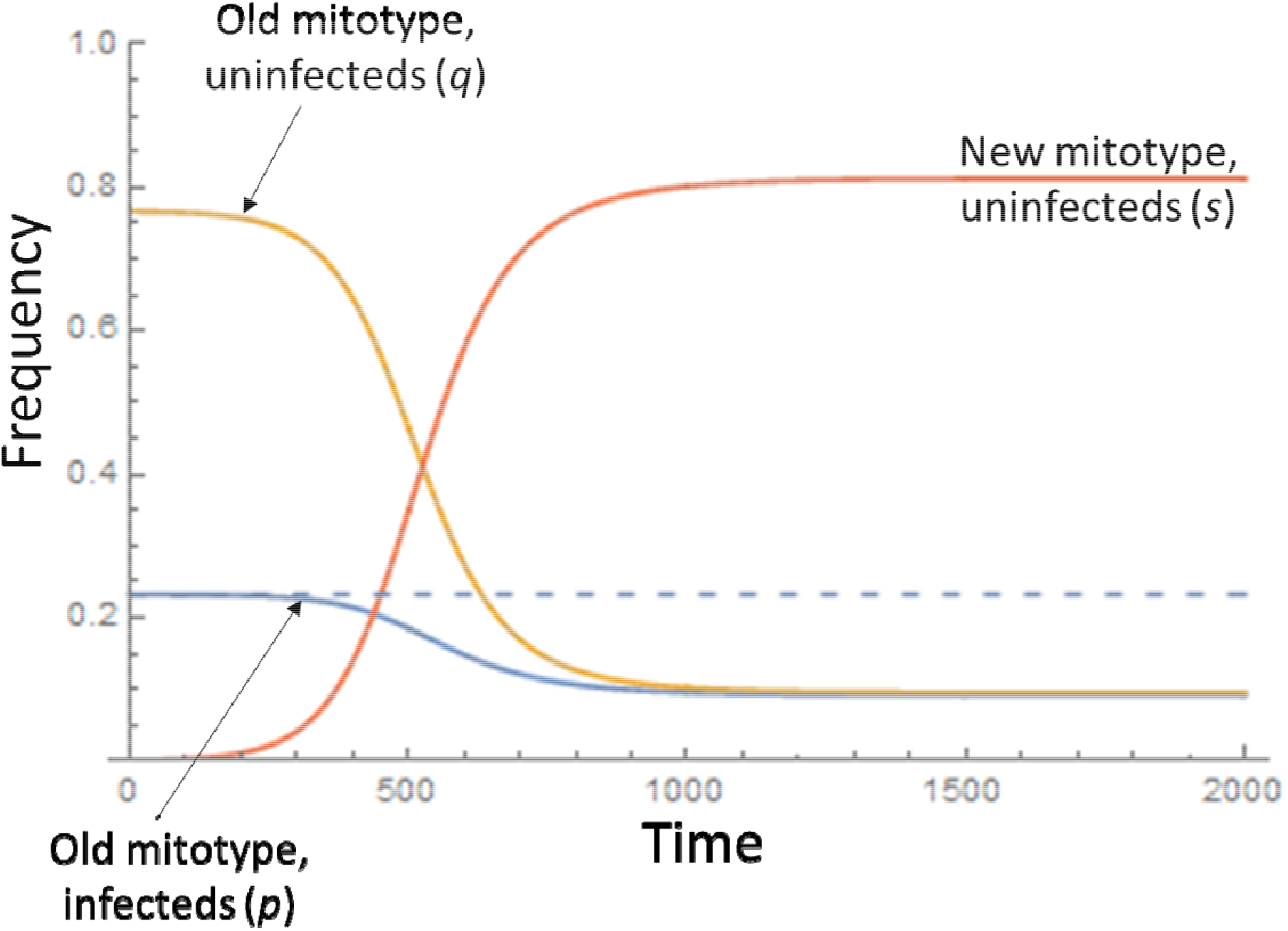
Dynamics of a beneficial mitochondrial type (*t* = 0.02) arising in an uninfected individual for a symbiont with frequency dependent benefit (*b* = 0.1, *c* = 0.05, μ = 0.02). Blue line – frequency of symbiont with ancestral mitochdondrial type; red line – frequency of uninfected individuals with new mitotype, orange, uninfected individuals with ancestral mitotype.

For the special case of no segregational loss (μ = 0), a novel advantageous mitotype that arises in the uninfected portion of the population rises in frequency and fixes within that section of the population. The symbiont is either depressed in frequency (if *t* < b – *c*) or is driven from the population (if *t* > *b* – *c*).

### iii) Facultative heritable symbionts whose benefit increases as they become more common

We modelled the fate of a novel advantageous mitotype in a population carrying a symbiont maintained by cytoplasmic incompatibility (CI). Cytoplasmic incompatibility is a positively frequency dependent trait, in which the zygotes produced by infected fathers and uninfected mothers die at a rate *h*, such that a fraction (1 – *h*) survive. As the frequency of infection in males increases, so does the rate at which uninfected cytotypes become inviable. We examined the fate of mitochondrial mutants that arose in a population at the upper (stable) equilibrium for the symbiont (see Appendix iii).

Taking *t* = 0.02 as the upper limit of possible levels of benefit to a mitochondrial mutation, it was unlikely for a novel mitotype arising in an uninfected individual to invade the population for any level of CI above *h*=0.05, which is very weak CI (Figure 5). Beneficial mitochondrial mutations arising in the symbiont-infected lineage spread to fixation (providing the benefit *t* > 1), moving into the uninfected section of the population following segregational loss of the symbiont (not shown).

**Figure 5:**
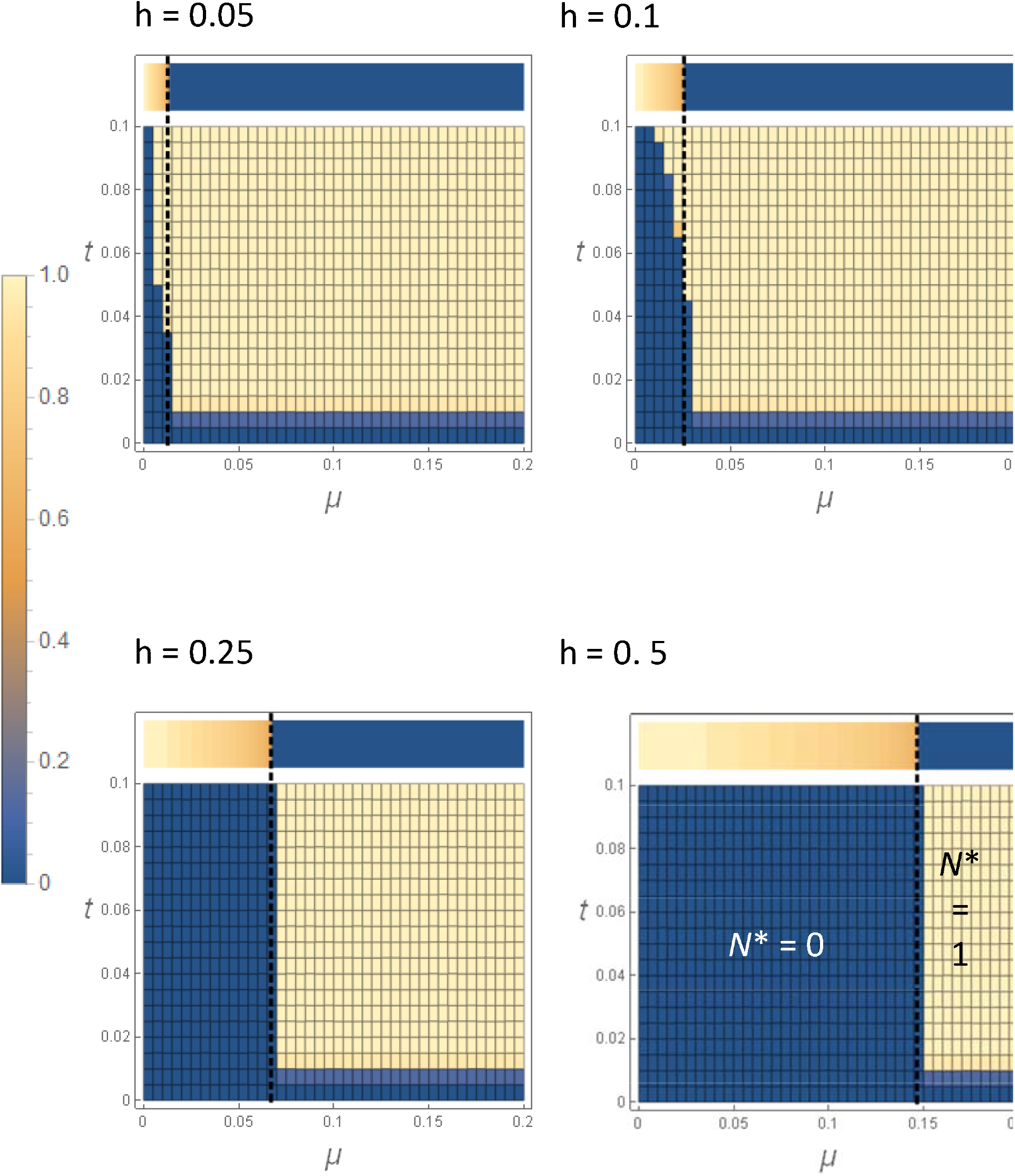
Conditions for invasion of a beneficial mitochondrial mutant arising into a symbiont-uninfected individual for different strengths of cytoplasmic incompatibility, *h*. The colours in the plots, relating to the scale at the left, show the overall equilibrium prevalence of the invading mitotype (*N*\* = *r*\* + s* yellow = fixation, blue = absent); the new mitotype invades and replaces the resident in the parameter space in yellow. X-axis is the rate of segregational loss (*μ*), and Y axis advantage of novel mitotype (*t*). The heat bar across the top of each plot is the frequency of the CI symbiont initially, before mitotype invasion (again, yellow = fixation, blue = absent). The symbiont cannot persist to the right of the dashed vertical line given by 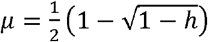.

## Discussion

Heritable microbial symbionts and mitochondria are both inherited through the female line, connecting the dynamics of the two elements. The relationship between microbial symbionts and the mitochondria with which they are coinherited has been well explored in terms of the impact of selection on symbionts driving the evolution of mitochondria (4, 17). Multiple studies have observed symbiont spread driving up the frequency of the associated mitotype. These effects are sufficiently strong that heritable symbionts can drive mtDNA across species boundaries following rare hybridization events (6,18). Historically, evolutionary biologists have viewed mtDNA as a neutral marker, useful for reflecting patterns of population demography and gene flow. However, both molecular evolution analyses and directed study of phenotype indicate mtDNA commonly undergoes positive selection (7–9). These data imply that selection on mtDNA may impact symbiont dynamics, which we tested here. Our analyses centre on the impact of novel, beneficial mutant mitotypes under directional selection on the dynamics of an existing facultative heritable symbiont.

Our analyses indicate that positive selection on mitochondria may drive symbionts extinct when the mutation creating a beneficial mitotype arises in an individual that does not carry the symbiont. The likelihood of this occurring is highest where the symbiont is at low frequency in the population. There are two reasons for such deterministic extinction being most commonly seen in low prevalence infection, beside simple stochastic fadeout at low frequencies. First, positive selection on mtDNA types will only exclude a symbiont where the mutation occurs initially in an uninfected individual, a situation that balance of numbers makes most likely for low prevalence symbiont infections. Second, symbiont frequency is determined by a balance of drive (the increase in the number of surviving daughters produced by an infected female compared to an uninfected female) and segregational loss (failure of progeny to inherit). The novel mitotype invades if carriers, on average, leave more infected daughters than left by the symbiont infected ones (carrying the ancestral mtDNA). For selective coefficients on the mtDNA of up to 2%, this only occurs for low prevalence symbionts.

Low prevalence symbiont infections are thought to be common. Historical studies of well understood systems where the symbiont has a strong phenotype, such as the *A. bipunctata* – *Rickettsia* interaction, *Drosophila bifasciata* – *Wolbachia* and *Drosophila* – *S. poulsonii* interaction, indicate 7%, 6% and 1-3% of individuals are infected in these species respectively (19–21). PCR screens also reveal low prevalence infections are common (22), and statistical models of screen data predict a majority of heritable symbiont infections occur rarely within a species (23). Whilst the data in screen studies are less robust (a low false positive rate in PCR assay likely inflates estimates of low prevalence infections), it is nevertheless clear that low prevalence infections are common. In conjunction with the belief that mtDNA is not a neutral marker, these data support the process modelled here occurring sufficiently commonly to represent an important contributor to symbiont turnover.

The lack of shared facultative heritable symbionts in sibling species pairs clearly indicates that loss of facultative heritable microbes from a population occurs commonly. However, the processes that produce symbiont loss are poorly understood. Mitochondrial evolution represents a novel potential driver of this process and is conceptually equivalent to observations of one symbiont replacing another, a process that has been observed in natural populations (24). The mitochondrial displacement process we have described will act alongside other factors, such as loss of the drive phenotype through suppression by the host (25), loss of the advantage to the phenotype due to ecological change (e.g. loss of a natural enemy for a protective symbiont) or mutational degradation of redundant symbiont phenotype (26).

The association of mitotype and symbionts is also expected to impact symbiont diversity. Where positively selected mitotypes arise in a symbiont infected individual, these episodes of selection result in the fixation of the symbiont strain originally associated with the beneficial mitotype. Thus, just as the level of mitochondrial diversity may be a function of the history of selection on the symbiont, so symbiont diversity may reflect historical selection on mtDNA. Notably, even weakly selected mitochondria are expected to impact symbiont diversity where these arise initially in an individual carrying a symbiont. Thus, interpretation of a population in which there is a lack of symbiont/mtDNA diversity requires more nuance than the traditional view in which symbiont evolutionary processes dominate the evolution of a neutral marker. A lack of variation in mtDNA and symbionts may reflect a selective sweep event on the symbiont, or a sweep event on the mtDNA.

Finally, our results indicate a further way in which symbionts may affect mitochondrial evolution. Weakly advantageous novel mitotypes will only spread where the mutation occurs in a symbiont-infected lineage. Where symbionts are at prevalence values <20%, these mutations are likely to occur in uninfected individuals and then fail to spread, or in the case of a symbiont with frequency dependent benefit, spread but be restricted to the uninfected fraction of the population. Thus, the presence of heritable symbionts may increase the time before beneficial mitotypes spread, as weakly beneficial mutations can only invade if they occur initially in symbiont infected individuals. This process is conceptual similar to background selection for linked nuclear loci, where weakly beneficial mutations do not spread if they initially occur near deleterious mutations (27).

The conclusions of our models depend on the assumption that symbiont and mitochondria are co-transmitted through maternal inheritance. This conclusion is validated in part by the many observations of sweeps of mtDNA associated with *Wolbachia* spread. Nevertheless, co-transmission may be disrupted through rare paternal mtDNA transmission (28), or intraspecific infectious transmission of the symbiont. Phylogenomic analysis comparing mtDNA and symbiont phylogenies indicate a high level of concordance within species, although some discordance is commonly observed (29–31). In general, the low level observed is unlikely to disrupt the processes we have been modelling, as the processes we are examining are occurring on short population genetic timescales (compared to the long timescales captured in phylogenomic analyses). Nevertheless, there are sporadic cases where symbiont infectious transmission is detectable in the timescale of laboratory experiments, particularly in parasitic wasp hosts (32, 33), and some species examined show no concordance of mtDNA and symbiont phylogeny (31). In these cases, our model assumptions are likely to be sufficiently compromised that the process we have predicted would not be manifested.

In conclusion, the coinheritance of mtDNA and maternally inherited symbionts means that selection on mtDNA can alter the dynamics and diversity of the symbiont, both in terms of reducing symbiont diversity within the population and driving the loss of low prevalence facultative symbionts. In female heterogametic taxa, these processes would extend to the associated W chromosome (34), and therefore be even more widespread. The evolution of cytoplasmically inherited component of the gene requires consideration of selection occurring in any and all components.

## Acknowledgements

We thank the UKRI for funding (grant NE/S012346/1).

## Appendix

### i) Facultative heritable symbionts with a fixed benefit

We assume a weakly-beneficial symbiont, such that offspring of infected mothers receive a net fitness benefit (1+B); this benefit is a composite of the fixed benefit supplied and the fixed costs of symbiont carriage. The symbiont is assumed to have vertical transmission efficiency (1 - μ), such that a proportion μ of offspring born to infected mothers do not carry the symbiont (segregational loss). As stated above, the new mitotype is assumed to confer a selective benefit *t* to individuals that carry it. The relative numbers of viable offspring born to each maternal type are given in Table 1. The change in frequencies of each demographic group from generation *T* to *T*+1 are then given by the recursion equations:

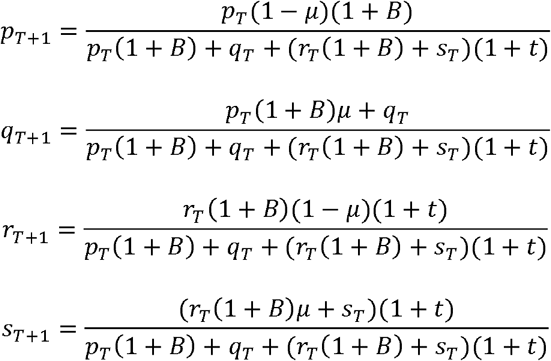

**Table 1.** Summary of relative numbers of viable offspring (generation *T*+1) born to each maternal cytotype (generation *T*) for the weakly beneficially symbiont model. *U* and *I* refer to cytotypes uninfected or infected respectively by the symbiont, and *R* and *N* refer to cytotypes carrying either the resident or new mitotypes respectively.

| Maternal cytotype | Freq. | $I_{R,T+1}$ | $U_{R,T+1}$ | $I_{N,T+1}$ | $U_{N,T+1}$ |
| --- | --- | --- | --- | --- | --- |
| $I_{R,T}$ | $p_T$ | $(1-\mu)(1+B)$ | $(1+B)\mu$ | 0 | 0 |
| $U_{R,T}$ | $q_T$ | 0 | 1 | 0 | 0 |
| $I_{N,T}$ | $r_T$ | 0 | 0 | $(1+B)(1-\mu)(1+t)$ | $(1+B)\mu(1+t)$ |
| $U_{N,T}$ | $s_T$ | 0 | 0 | 0 | $(1+t)$ |

First we establish the conditions for symbiont to persist in the presence of the resident mitotype alone (*r* = *s* = 0). Here the system can be described by the single equation (since *q* = 1 – *p*):

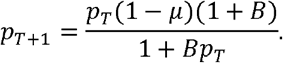

There are 2 equilibria to this system: *p*\* = 0 and 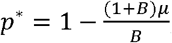; the latter of which is stable if 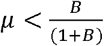, being the criterion for symbiont persistence (as shown by the vertical line in Fig 2; the heatmap in Fig 2 corresponds to the value of *p*\*).

We now establish the stability criteria of the full system to determine under what conditions the new mitotype can invade, and the consequences for the symbiont. There are 4 equilibria (presented in the order {*p*\*,*q*\*,*r*\*,*s*\*}):

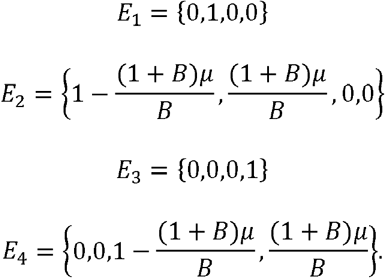

An equilibrium is stable if the magnitude of all the eigenvalues calculated from the Jacobian matrix evaluated at that equilibrium are less than 1. Both *E_1_* and *E_2_*, involving the persistence of the resident mitotype, have eigenvalues (1 + *t*) and so are always unstable, assuming *t* > 0. Hence, assuming the new mitotype confers a non-zero selective advantage over the resident mitotype, it should invade and replace the resident. Indeed, the equilibrium *E_4_* (symbiont persistence with the new mitotype alone) is always stable providing *t* > 0 and the standard criterion for symbiont persistence, 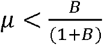, is upheld. So, providing the new mitotype arises in a symbiont-infected individual, the new mitotype will sweep in, replacing the resident mitotype, and bringing the symbiont with it (Fig 1).

However, the eigenvalues for *E_3_* (the new mitotype in the absence of the symbiont) include 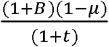. This eigenvalue is greater than 1, and hence this equilibrium is unstable, if t < *B*(1 – *μ*) – *μ* (shown by the diagonal line in Fig 2). Hence in this region, the system involving the new mitotype in the absence of the symbiont is not stable. In other words, if t < *B*(1 – *μ*) – *μ*, then the new mitotype cannot invade and replace the resident if it initially arises in an uninfected individual; the state *E_4_*, containing the symbiont, would be stable in this region of parameter space, but it can never be attained if the new mitotype is restricted to the uninfected population.

### ii) Facultative heritable symbionts whose benefit declines as they become more common

Here we model a symbiont whose benefit declines with its frequency, and for simplicity we assume a linear relationship between the benefit *B* and overall symbiont frequency (*p* + *r*) of the form *B_T_* = *b*(1 – (*p_T_* + *r_T_*)), where *b* is the maximum benefit when the symbiont is vanishingly rare. We also include a cost, *c*, to symbiont carriage, for example through a reduction in host fecundity. As before we assume a rate of segregational loss of the symbiont, μ. The numbers of viable offspring born to each maternal type are given in Table 2, and the change in frequencies of each demographic group from generation *T* to *T*+1 are then given by the recursion equations:

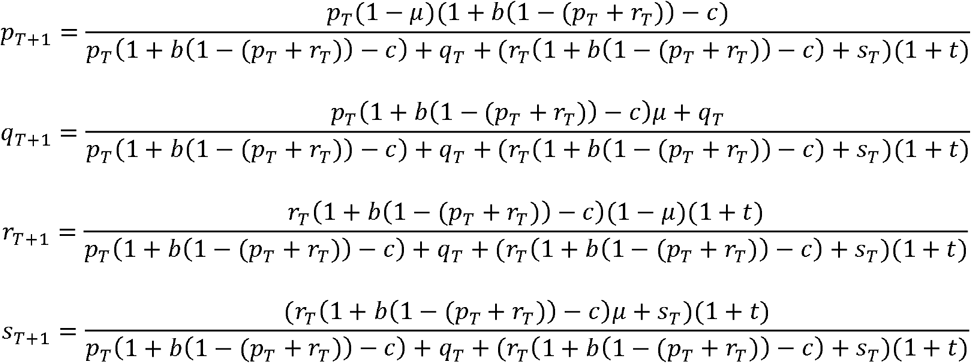

**Table 2.** Summary of relative numbers of viable offspring (generation *T*+1) born to each maternal type (generation *T*) for the negative frequency-dependent benefit model. *P_T_*(= *p_T_* + *r_T_*) is the overall frequency of the symbiont in the population, regardless of mitotype, in generation *T*.

| Maternal<br>cytotype | Freq. | $I_{R,T+1}$ | $U_{R,T+1}$ | $I_{N,T+1}$ | $U_{N,T+1}$ |
| --- | --- | --- | --- | --- | --- |
| $I_{R,T}$ | $p_T$ | $(1-\mu)(1+b(1-P_T)-c)$ | $(1+b(1-P_T)-c)\mu$ | 0 | 0 |
| $U_{R,T}$ | $q_T$ | 0 | 1 | 0 | 0 |
| $I_{N,T}$ | $r_T$ | 0 | 0 | $(1+b(1-P_T)-c)(1-\mu)(1+t)$ | $(1+b(1-P_T)-c)\mu(1+t)$ |
| $U_{N,T}$ | $s_T$ | 0 | 0 | 0 | $(1+t)$ |

As before we establish the conditions for symbiont persistence in the presence of the resident mitotype alone (*r* = *s* = 0). Here the system can be described by the equation:

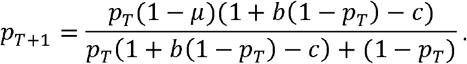

There are 2 equilibria to this system: *p*\* = 0 and 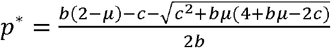, which is stable if 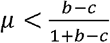, the criterion for symbiont persistence (vertical line in Fig 3; again the heatmap in Fig 3 corresponds to the above value of *p*\*).

We now establish the stability criteria of the full system. There are now 8 equilibria, corresponding to different combinations of symbiont presence or absence, in the presence of either the resident or the new mitotype, and some of the equilibria involve multiple polynomial roots. The biologically-relevant ones (presented in the order {*p*\*, *q*\*,*r*\*,*s*} are:

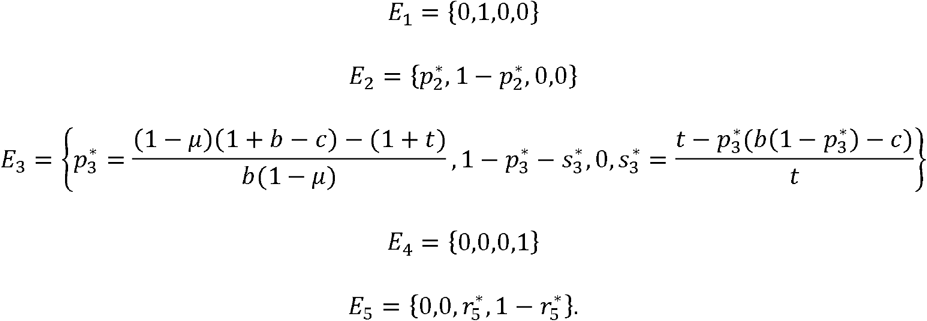

where 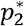 and 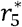 are both given by the expression for the equilibrium symbiont frequency, defined above, just in one case (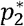) it occurs with the resident mitotype on its own, and in the other case (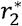) it occurs with the new mitotype on its own. Note that equilibrium *E_3_* involves coexistence of both mitotypes (but see below for explanation of this state).

A full analytical assessment of all eigenvalues of this system is not possible, but a combination of analysis of those states that are analytically tractable and numerical analyses reveals the following points. First, all equilibria involving the resident mitotype (states *E_1_* – *E_3_*) have (1 + *t*) as one of their eigenvalues, so these states are always unstable, providing the new mitotype has some selection advantage (*t* > 0). Second, state *E_5_*, involving the new mitotype alone carrying the symbiont, is always stable providing *t* > 1 and the general criterion for symbiont persistence 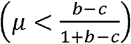 is upheld. Hence the new mitotype should always be able to sweep in, replace the resident mitotype, and carry the symbiont with it (which should achieve the same equilibrium prevalence as in the resident). That is what happens if the new mitotype arises in a symbiont-infected individual; however, if the new mitotype arises in an uninfected individual, state *E_5_* cannot be achieved since the symbiont cannot jump between maternal lines. In that case we may get different outcomes from among the remaining (unstable) states.

In particular, there are 2 key boundaries that separate the different outcomes when the new mitotype arises in an uninfected individual (Fig 3, diagonal and humped lines). The diagonal line, obtained from analysis of the eigenvalues of state *E_4_*, is given by the expression *t* = (1 + *b* – *c*)(1 – *μ*) – 1. Above this line the new mitotype invades and replaces the resident but, since it arises in an uninfected individual, it cannot be infected by the symbiont, and so the system stays at *E_4_*, unable to switch to state *E_5_* due to the inability of the symbiont to jump into the new mitotype. Below the diagonal line (*t* < (1 + *b* − *c*)(1 − *μ*) – 1) one of 2 outcomes can occur, depending on the second boundary (the humped line in Fig 3), obtained by setting the eigenvalue 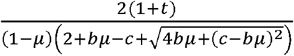 from state *E_2_* equal to 1 and solving for *t*, to give: 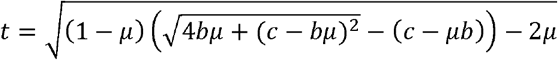. Above this line (but below the diagonal line) the new mitotype can invade, but it doesn’t replace the resident mitotype. Again, the new mitotype cannot carry the symbiont, which remains in the resident mitotype (i.e., the system remains at state *E_3_*, again unable to jump to *E_5_*). Notably the symbiont is reduced to a lower overall prevalence than previously, due to the reduction in frequency of the resident mitotype in which it occurs, as a result of (partial) invasion by the new mitotype. However, below the humped line the new mitotype cannot invade, and the system remains at *E_2_*, with the symbiont persisting in the resident mitotype at its original frequency.

### iii) Facultative heritable symbionts whose benefit increases as they become more common

Finally we consider a symbiont under positive frequency dependent selection, like one that confers cytoplasmic incompatibility (CI). In such a model it is assumed that crosses between uninfected females and infected males reduce offspring fitness by a factor (1 – *h*). As before we also assume a rate of segregational loss of the symbiont, μ. For simplicity we ignore potential costs of symbiont carriage, as at equilibrium these are vastly outweighed by benefits under positive frequency dependence.

Because we have to consider the effect of different male-female matings to account for CI, we now explicitly keep track of the frequencies of uninfected and infected males and females of each mitotype. The numbers of viable offspring born to each maternal-paternal combination are given in Table 3, and the overall changes in frequencies of each group from generation *T* to *T*+1 are:

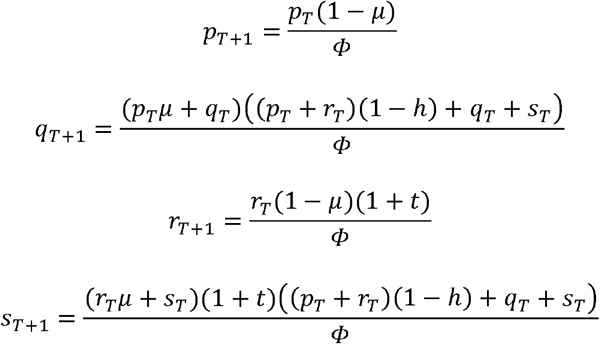

where *Φ* is the sum of all the offspring values in Table 3, weighted by their corresponding mating frequency.

**Table 3.**
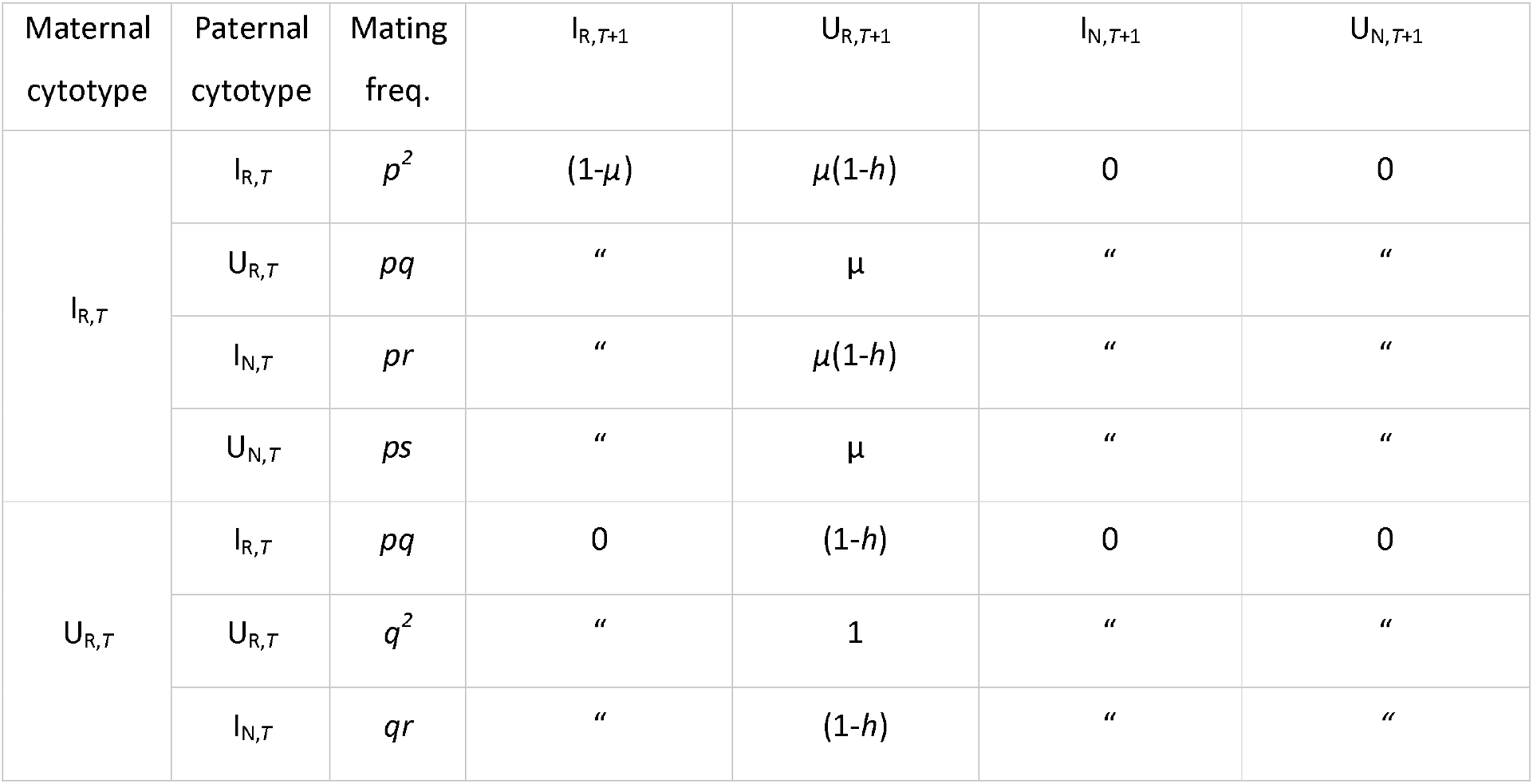
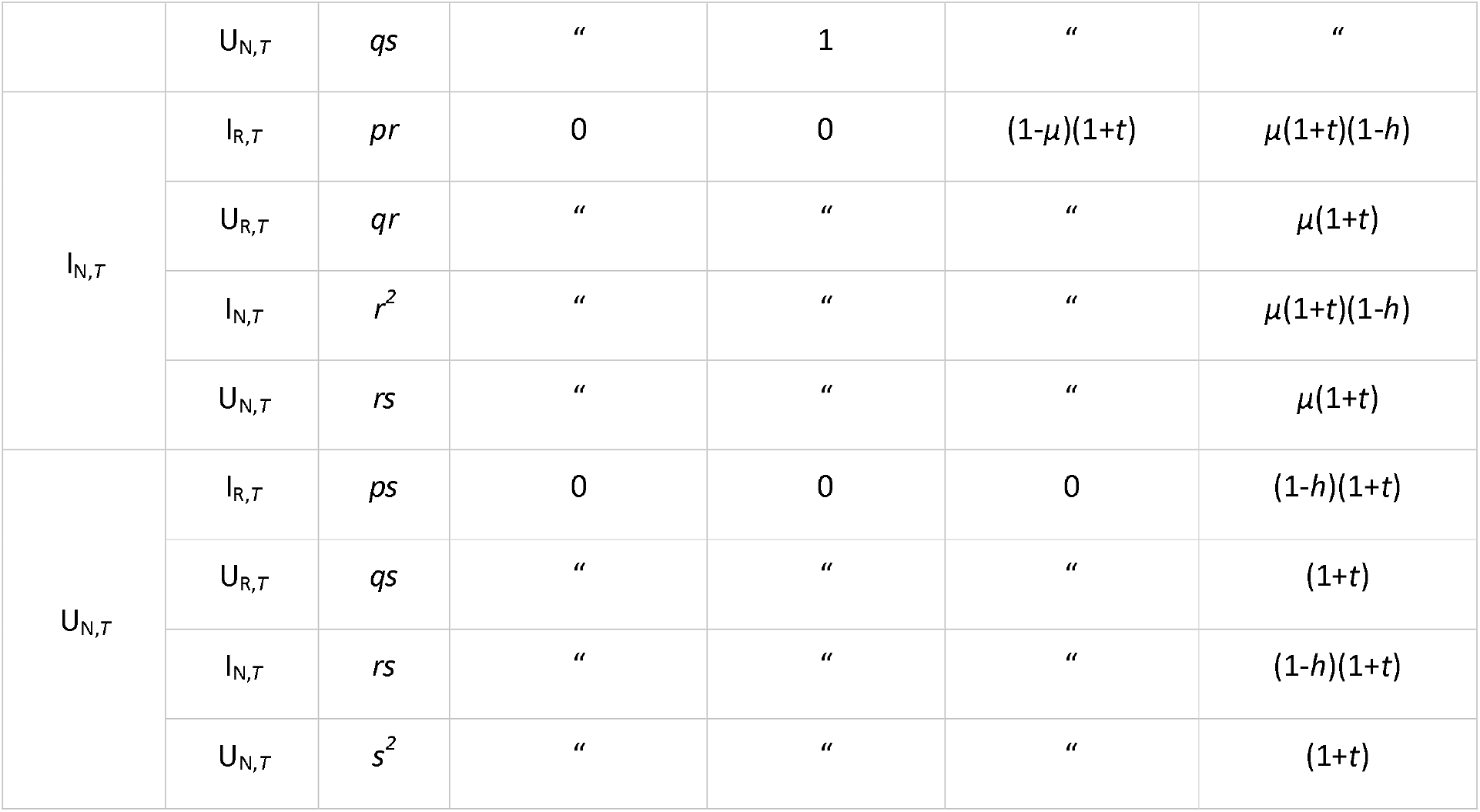
Summary of relative numbers of viable offspring (generation *T*+1) born to each mating combination (generation *T*) for the positive frequency-dependent (CI) model.

| Maternal<br>cytotype | Paternal<br>cytotype | Mating<br>freq. | $I_{R,T+1}$ | $U_{R,T+1}$ | $I_{N,T+1}$ | $U_{N,T+1}$ |
| --- | --- | --- | --- | --- | --- | --- |
| $I_{R,T}$ | $I_{R,T}$ | $p^2$ | $(1-\mu)$ | $\mu(1-h)$ | 0 | 0 |
| | $U_{R,T}$ | $pq$ | " | $\mu$ | " | " |
| | $I_{N,T}$ | $pr$ | " | $\mu(1-h)$ | " | " |
| | $U_{N,T}$ | $ps$ | " | $\mu$ | " | " |
| $U_{R,T}$ | $I_{R,T}$ | $pq$ | 0 | $(1-h)$ | 0 | 0 |
| | $U_{R,T}$ | $q^2$ | " | 1 | " | " |
| | $I_{N,T}$ | $qr$ | " | $(1-h)$ | " | " |
| | $U_{N,T}$ | $qs$ | $''$ | $1$ | $''$ | $''$ |
| $I_{N,T}$ | $I_{R,T}$ | $pr$ | $0$ | $0$ | $(1-\mu)(1+t)$ | $\mu(1+t)(1-h)$ |
| | $U_{R,T}$ | $qr$ | $''$ | $''$ | $''$ | $\mu(1+t)$ |
| | $I_{N,T}$ | $r^2$ | $''$ | $''$ | $''$ | $\mu(1+t)(1-h)$ |
| | $U_{N,T}$ | $rs$ | $''$ | $''$ | $''$ | $\mu(1+t)$ |
| $U_{N,T}$ | $I_{R,T}$ | $ps$ | $0$ | $0$ | $0$ | $(1-h)(1+t)$ |
| | $U_{R,T}$ | $qs$ | $''$ | $''$ | $''$ | $(1+t)$ |
| | $I_{N,T}$ | $rs$ | $''$ | $''$ | $''$ | $(1-h)(1+t)$ |
| | $U_{N,T}$ | $s^2$ | $''$ | $''$ | $''$ | $(1+t)$ |

Again we establish the conditions for symbiont persistent in the presence of the resident mitotype alone (*r* = *s* = 0). Here the system can be described by the equation:

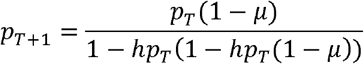

There are 3 equilibria to this system:

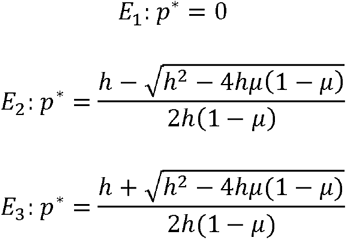

*E_3_* is the positive internal equilibrium for the symbiont, which is biologically feasible if 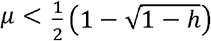, the expression for the vertical line in Fig 5 (the heatmap at the top of each plot in Fig 5 corresponds to the above value of *p*\* from *E_3_*). However, due to the positive frequency dependency inherent in the CI system, this criterion is not sufficient to guarantee symbiont persistence (e.g., Hoffman 1990). State *E_2_* is an unstable equilibrium, such that the initial frequency of the symbiont, *p*_0_, has to exceed that equilibrium value in order to reach the internal equilibrium *E_3_*; if not then the system heads to *E_1_* and the symbiont fails to persist. Unfortunately a mathematical analysis of the full system involving the invading mitotype is not possible – for that reason we resort to a numerical analysis based on simulations to generate the results shown in Fig 5, focussing on the conditions for the new mitotype to invade, when arising in the uninfected lineage.

